# Unifying the Identification of Biomedical Entities with the Bioregistry

**DOI:** 10.1101/2022.07.08.499378

**Authors:** Charles Tapley Hoyt, Meghan Balk, Tiffany J. Callahan, Daniel Domingo-Fernández, Melissa A. Haendel, Harshad B. Hegde, Daniel S. Himmelstein, Klas Karis, John Kunze, Tiago Lubiana, Nicolas Matentzoglu, Julie McMurry, Sierra Moxon, Christopher J. Mungall, Adriano Rutz, Deepak R. Unni, Egon Willighagen, Donald Winston, Benjamin M. Gyori

## Abstract

The standardized identification of biomedical entities is a cornerstone of interoperability, reuse, and data integration in the life sciences. Several registries have been developed to catalog resources maintaining identifiers for biomedical entities such as small molecules, proteins, cell lines, and clinical trials. However, existing registries have struggled to provide sufficient coverage and metadata standards that meet the evolving needs of modern life sciences researchers. Here, we introduce the Bioregistry, an integrative, open, community-driven metaregistry that synthesizes and substantially expands upon 23 existing registries. The Bioregistry addresses the need for a sustainable registry by leveraging public infrastructure and automation, and employing a progressive governance model centered around open code and open data to foster community contribution. The Bioregistry can be used to support the standardized annotation of data, models, ontologies, and scientific literature, thereby promoting their interoperability and reuse. The Bioregistry can be accessed through https://bioregistry.io and its source code and data are available under the MIT and CC0 Licenses at https://github.com/biopragmatics/bioregistry.

## 1 Introduction

One of the key challenges in creating and maintaining Findable, Accessible, Interoperable, and Reusable (FAIR)^1–3^ data in the life sciences is the standardized identification of entities ranging from chemicals, proteins, and diseases to patents and publications. These entities are typically curated in *identifier resources* (e.g., ontologies and databases) such as Chemical Entities of Biomedical Interest (ChEBI)^4^, UniProt^5^, and PubMed that assign to each entity a *local unique identifier* (i.e., accession number). Each resource defines an internally consistent pattern for its entities’ local unique identifiers, such as the combination of numbers and letters found in UniProt identifiers (e.g., P0DP23) or the simple numbers found in PubMed identifiers (e.g., 29175850). Uniform resource identifiers (URIs) (e.g., https://www.uniprot.org/uniprot/P0DP23) and compact uniform resource identifiers (CURIEs) (e.g., uniprot:P0DP23) have become the predominant syntaxes used in the life sciences for identifying entities that encode both the resource from which the entity originates and its local unique identifier^6^. URIs encode the resource with a *URI prefix* (e.g., https://www.uniprot.org/uniprot/) while CURIEs encode it with a *prefix* (e.g., uniprot).

However, even when using URIs and CURIEs, a number of challenges remain in establishing consistency and interoperability. Namely, several different incompatible URIs and CURIEs can be used to refer to the same entity. For example, the local unique identifier P0DP23 for the entry in UniProt^5^ about the Calmodulin-1 protein can be represented by at least seven distinct URIs and three distinct CURIEs (see Supplementary Tables 1 and 2 for details). This problem is compounded when attempting to integrate multiple resources, a cornerstone of modern computational life sciences. For example, genomic data from HGNC^7^ can not be readily integrated with biochemical reactions data from Rhea^8^ because HGNC uses the prefix ec-code and Rhea uses the prefix EC when referring to entities in the Enzyme Commission identifier resource^9^. Similarly, many biomedical resources construct local unique identifiers for the same entity in the Enzyme Commission identifier resource differently, e.g., 1.4 in IntEnz^10^, 1.4.-.- in the Gene Ontology (GO)^11^, and 1.4.* in ChEBI for *Oxidoreductases acting on the CH-NH2 group of donors*.

In order to standardize the usages of URIs and CURIEs and therefore enable their interoperability, a *registry* is needed containing canonical, validatable definitions of identifier resources that, for each resource, includes a prefix, a URI prefix, a local unique identifier pattern, and other associated metadata. Registries thus capture for each identifier resource how to construct, parse, and interchange canonical URIs and CURIEs. A registry can be used by external biomedical resources to standardize the way they reference entities (e.g., database cross-references appearing in ontologies) to promote integration with other resources, as well as by consumers to navigate prefixes and their associated metadata. Multiple registries ^5,11–30^ have been previously built for this purpose, but they each suffer from substantial gaps in their coverage of known resources and the metadata captured about these resources. They also lack interoperability among each others’ entries, for example, the National Center for Biotechnology Information Taxonomy Database (NCBITaxon)^31^ is prefixed as taxonomy in Identifiers.org^23^ and as NCBITAXON in BioPortal^16^.

These issues are exacerbated by shortcomings in existing registries’ governance and curation workflows, which impede their ability to stay current, trustworthy, and engage the community as the landscape of life science resources rapidly evolves. These issues include, that they 1) are built on private infrastructure within an institution; 2) are maintained by small, private groups that - due to under-funding - struggle to respond to requests; 3) lack adequate support for external contributions 4) are neither versioned nor archived. As an alternative to general-purpose registries, numerous projects (e.g., GO, Cellosaurus^17^, NCBI GenBank^24^) have created their own registries, however, these each only cover identifier resources relevant for the given project, and use standards that are only internally consistent to the project. Finally, several services act as registries but are by design limited in scope to include a selected set of resources and provide incomplete metadata necessary to promote the standardization of references. These include the Open Biomedical Ontologies (OBO) Foundry^25^, Ontology Lookup Service (OLS)^26^, and BioPortal^16^. A detailed survey of the governance and maintenance models for existing registries can be found in Supplementary Table 3. Overall, the content of any one registry does not reflect the evolving landscape of biomedical resources and thus satisfy user needs.

To overcome the limitations of existing registries, a new approach for building a biomedical registry is necessary, which ideally fulfills the three requirements: 1) integrative, 2) open, and 3) community-driven. First, an *integrative* registry re-uses, improves, and extends on existing registries. Given that existing registries define conflicting standards (i.e., assign conflicting prefixes, URI prefixes, or other metadata for the same identifier resource) and therefore lack interoperability, this necessitates alignment and harmonization among resources in each registry. Second, an *open* registry makes its underlying data and associated code available under permissive licenses in a public, version-controlled repository, and relies on free, publicly available infrastructure for semi-automated quality control and deployment. Third, a *community-driven* registry solicits contributions from community members and provides an appropriate technical platform and governance structure to support this. This technical platform needs to support discussion and feedback as well as quality assurance workflows tightly coupled to the underlying open data, code, and infrastructure. Overall, these properties are expected to promote the sustainability and longevity of the registry.

To address these limitations, we introduce the Bioregistry: an integrative, open, and community-driven metaregistry. The Bioregistry integrates content from existing registries, semi-automatically identifies equivalences between records in existing registries, and resolves conflicts between them using a novel workflow. The result of this alignment makes the Bioregistry a *metaregistry* (i.e., a registry of registries) which, for each resource, maintains cross-registry mappings to serve as an interoperability layer between conflicting standards. The Bioregistry also includes substantial manual curation for resources not appearing in any pre-existing registry, and additional curation extends and improves on the metadata associated with resources in these registries with curation practices heavily influenced by the most similar existing registries (e.g., Identifiers.org and Prefix Commons). As a result of this process, the Bioregistry expands substantially on the content of each individual pre-existing registry (e.g., 81% over Identifiers.org) as well as all aligned registries combined (see Table 2 on alignment) and incorporating feedback from the maintainers and developers of existing registries. The Bioregistry also provides a higher granularity data model compared to any existing registry thereby better supporting integration. The Bioregistry is built using open source code, open data, and leverages public infrastructure and automation to support its maintenance and extension. Further, it has well-defined contribution guidelines and a multi-institutional governance model that enables contributions directly from the broader community to support the project’s longevity.

**Table 1.**
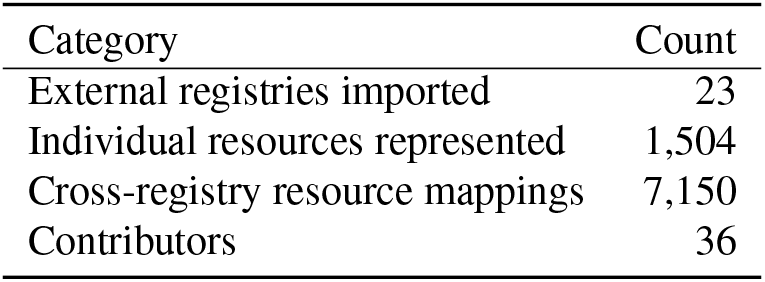
Overview statistics of the Bioregistry version 0.5.132 (2022-10-17).

**Table 2.**
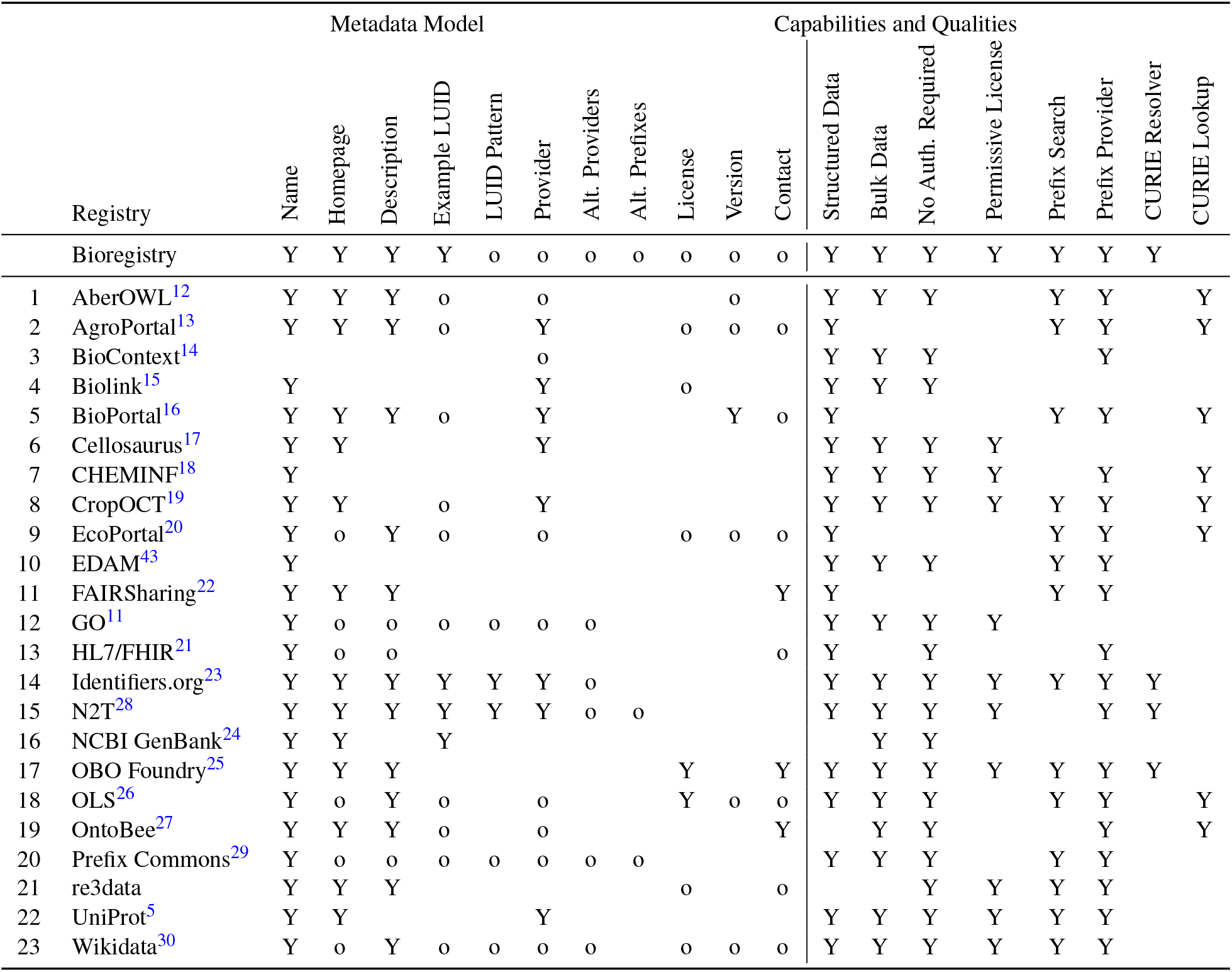
An overview of the data models, capabilities, and qualities of the 23 external registries covering biomedical ontologies, controlled vocabularies, and databases imported by the Bioregistry. A “Y” means the field is required. A “o” means it is part of the schema, but not required or incomplete on some entries. A blank cell means that it is not part of the metadata schema. Two glossaries describing the elements in the data model (left) and capabilities and qualities (right) appearing as column headers in this table can be found respectively in Supplementary Sections 3.1 and 8.1.

The Bioregistry (0.5.132) integrates and aligns content previously curated in 23 external registries, and in total, contains 1,504 individual records. These records extend on each prior registry (as compared to e.g., 838 records in BioContext^14^ and 774 in Identifiers.org^23^), as well as all aligned registries combined: 192 of the Bioregistry’s 1,504 records are novel, i.e. they do not appear in any existing registry. The Bioregistry also adds novel curated metadata for 969 of the remaining 1,312 records (74% of all records). A summary of the content captured in Bioregistry is provided in Table 1. We provide detailed metrics and comparison to other resources in the Results section.

The Bioregistry has already been integrated with several projects aimed at data integration, knowledge assembly, and semantically annotated publications, described in section 3 on use cases and integrations. Current integrations of Bioregistry include BridgeDb^32^, PheKnowLator^33^, Manubot^34^, Biomappings^35^, SSSOM^36^, INDRA^37^, and the OBO Foundry^25^.

The Bioregistry is available through an interactive web portal (https://bioregistry.io), an OpenAPI-documented web service (https://bioregistry.io/apidocs), a Python software package (https://pypi.org/project/bioregistry), and a Docker image (https://hub.docker.com/r/biopragmatics/bioregistry). All underlying data, code, and governance documentation are accessible through GitHub (https://github.com/biopragmatics/bioregistry) under the MIT and CC0 licenses and archived on Zenodo^38^.

## 2 Results

### Bioregistry Data Model

The Bioregistry uses a granular and extensible data model to represent records of identifier resources. Required fields for each record include a canonical prefix, a human-readable label, a homepage, and a description. The data model also allows for multiple optional fields including the license, version, prefix synonyms for the resource, and capturing whether the resource is deprecated or proprietary. Each record can further include an example local unique identifier, a regular expression pattern for validating local unique identifiers, and a URI format string for constructing URIs from local unique identifiers. Records describing ontologies can include optional download links for associated OBO, OWL, and OBO Graph JSON artifacts. To keep contributions traceable and provide attribution, each record captures the submitter and reviewer who contributed to the entry. Records can also be grouped into collections for better contextualization, such as prefixes useful for the Semantic Web (e.g., DC, FOAF, RDF, RDFS).

A comparison between the data model and various properties of the Bioregistry and external registries in Table 2 demonstrates the heterogeneity of metadata standards in external registries and the flexibility of the Bioregistry to represent more granular metadata. For example, this enables the Bioregistry to represent deprecated and obsolete records for posterity, such as hgnc.genefamily (the HGNC Gene Family resource^39^) which was replaced by hgnc.genegroup, and casspc (Eschmeyer’s Catalog of Fishes^40^) which is used by the Teleost Taxonomy Ontology^41^ and Vertebrate Taxonomy Ontology^42^ but was itself never published.

Importantly, the Bioregistry captures not only individual resource records but also semantic relationships between records (including e.g., *depends on*, which asserts that one resource reuses terms from another, such as GO depends on ChEBI, and mappings of each resource to external registries where it appears. These additional relations constitute the Bioregistry’s *metaregistry*, a term meant to represent the fact that it creates links among previously incompatible resources through a set of cross-registry mappings.

The Bioregistry data model is described in further detail in Supplementary Section 3.

### Integration and harmonization of existing registries

The Bioregistry imports records from the 23 external registries described in Table 2. We divided registries into distinct groups to which we applied different import policies ranging from registries imported entirely to ones from which metadata for only select records are imported (see Methods). The Bioregistry uses a multi-stage process in which registries are sequentially imported such that a record from a given registry is either 1) aligned with an existing Bioregistry record and a cross-registry mapping is created 2) added as a new record, or 3) set aside for manual curation (see Methods). The key challenge in this import process is aligning (i.e., finding equivalences between) records since the external registries’ records are partially overlapping but inconsistent. This inconsistency stems from high heterogeneity among existing registries in the usage of capitalization (e.g., go vs. GO), punctuation (e.g. ec-code vs eccode), abbreviations (e.g., flybase vs. fb), and even different vocabulary (e.g., intenz vs. eccode) to represent the same resource, which results in fragmentation and lack of interoperability. A novel contribution of the Bioregistry is that it explicitly represents the results of its alignment procedure as equivalence mappings between its own records and records in external registries. This constitutes a network of cross-registry mappings (with Bioregistry as the hub), creating a metaregistry. The Bioregistry’s alignment procedure recovers a total of 7,150 such cross-registry mappings thereby connecting resources across existing registries that were previously disconnected. Importantly, the Bioregistry does not import nor redistribute the resources described by external registries, which themselves may have restrictive licenses (e.g., ICD-10). Further, rather than pushing enriched metadata back to the existing registries, the Bioregistry instead makes its content available under a permissive license in many formats (described in Figure 2) that can be individually processed by each external registry’s maintainers to best suit their needs.

**Figure 1.**
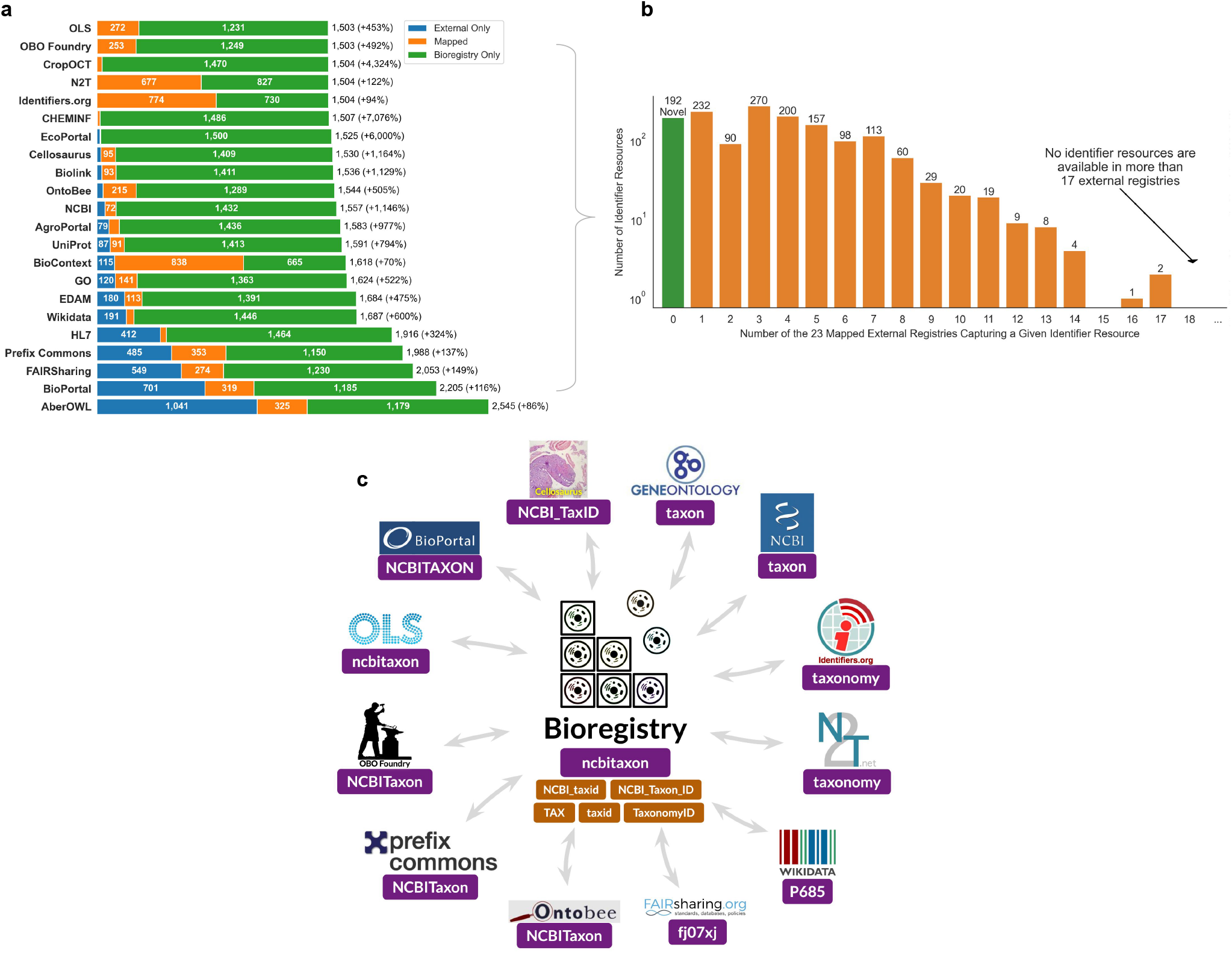
**A)** Summary of the pairwise overlap (in horizontal orange bars) between the prefixes in the Bioregistry and its integrated external registries. The horizontal blue bars show records that could not be automatically aligned and the horizontal green bars represent additional prefixes available in the Bioregistry but not the external resource. The absolute number of records in the union of the external registry with the Bioregistry (accounting for known overlaps) are shown on the right as well as the percentage relative gain introduced by the Bioregistry in parentheses. A large orange section corresponds to high content reuse while a large blue section corresponds to either high novelty of content in the external registry or high potential for semi-automated import into the Bioregistry. Counts on sections of these bar plots representing less than 70 prefixes are omitted. **B)** A histogram of how many cross references each entry in the Bioregistry has to external registries. The green bar highlights the prefixes with no cross references that only appear in the Bioregistry. **C)** A schematic diagram depicting the Bioregistry as an interoperability layer between external registries. Using the NCBI Taxonomy identifier resource as an example, prefixes used for this resource in external registries that the Bioregistry aligns are shown in purple boxes. Additional synonyms for this resource curated in the Bioregistry are shown in orange boxes. The components of this figure are regenerated daily with GitHub Actions and stored in https://github.com/bioregistry/bioregistry/tree/main/docs/img.

**Figure 2.**
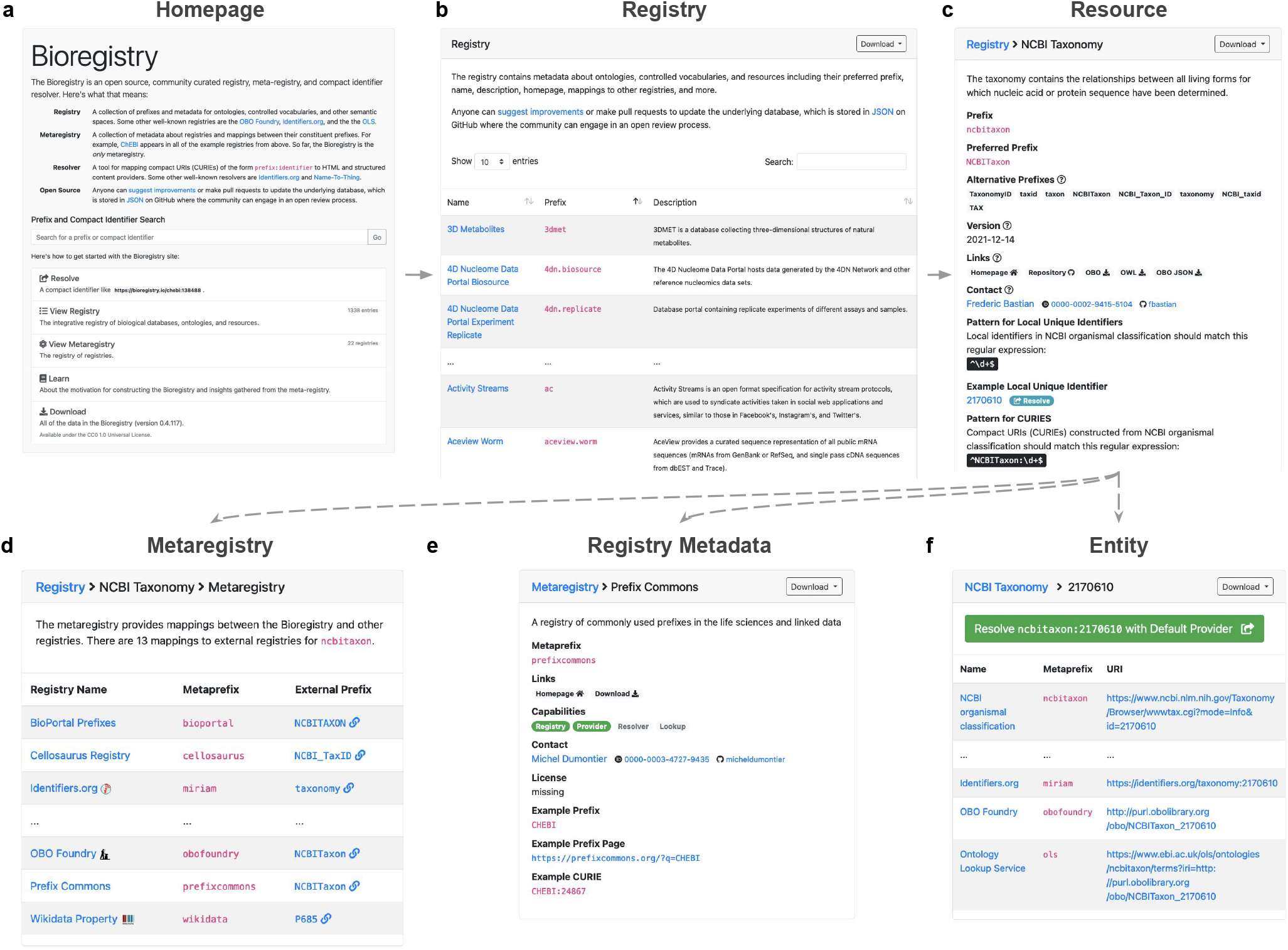
Website Screenshots. **A)** The homepage of https://bioregistry.io prominently features a combine prefix search and CURIE resolution box along with links to all of the components of the site. **B)** The full registry of prefixes, resource names, and descriptions can be viewed and full text search performed. **C)** Each prefix page shows metadata about the corresponding resource, its identifiers, and serves as a hub for additional functionality in D, E, and F. **D)** The prefix page additionally includes the metaregistry’s cross-registry mappings from the prefix to external registries’ prefixes. **E)** Each external registry page shows metadata and the capability list of external resources. **F)** a sample identifier demonstrates all of the providers that can be resolved.

We additionally curated 423 synonyms used when referring to identifier resources outside registries such as in OBO Foundry ontology database cross-references. These synonyms support the registry alignment workflow and broaden the ability of Bioregistry to standardize references to identifier resources beyond just external registries.

We investigated how the content of the integrated Bioregistry compares to each individual external registry that it imports and aligns (Figure 1A). It covers several registries (BioContext^14^, CHEMINF^18^, Crop Ontology Curation Tool (CropOCT)^19^, OBO Foundry^25^, OLS, Name-to-Thing (N2T)^28^, Identifiers.org^23^) almost entirely (over 85% of the external registries’ records are mapped to a Bioregistry record) while significantly expanding on the content of each of them from a minimum of +94% for Identifiers.org to a maximum of +7,076% for CHEMINF. The Bioregistry is able to align a smaller proportion of records in external registries such as FAIRSharing^22^ (33.3%), BioPortal^16^ (31.3%), Aber-OWL^12^ (23.8%), and Wikidata (20.7%) due to several characteristics of each registry. For example, many records in FAIRSharing^22^ do not refer to identifier resources, Wikidata contains many records lacking a biological scope, and Aber-OWL and BioPortal contain ontologies of heterogeneous quality which are queued for on-demand inclusion rather than automated ingestion into in the Bioregistry (see Methods). Despite lower coverage of their entries, the Bioregistry still substantially expands on these external registries between a minimum of +86% for Aber-OWL to a maximum of +6,000% for EcoPortal^20^.

We then investigated the frequency of appearance of each identifier resource in multiple registries (Figure 1B). We found that only 7 resources appeared in more than 13 of the 23 external registries (including well-known resources such as GO, ChEBI, and NCBITaxon), and no resource appeared in more than 17, further illustrating the fragmented state of existing registries and the benefits of an integrative registry in having improved coverage. Further, the Bioregistry contains 192 novel prefixes not available in any other registry (Figure 1B, green bar) that were curated for a diverse set of reasons. For example, we systematically reviewed cross-references in OBO Foundry ontologies, found a number of prefixes referring to resources for which no external registry contained an entry, and added them to the Bioregistry. Several novel prefixes were suggested by external contributors who were themselves the maintainers of the corresponding resource. Another subset of these entries were added by members of the community who encountered them and were then motivated to create an entry for it in the Bioregistry.

Next, we examined NCBITaxon, one of the resources that appears in the largest number of existing registries. This identifier resource appears in 17 external registries under 10 different prefixes including taxon, taxonomy, and NCBITaxon (Figure 1C, not all shown). In addition, the Bioregistry curates 9 prefix synonyms (e.g., NCBI_Taxon_id, uniprot.taxonomy, NCBI_taxid) that appear in various non-registry biomedical resources, demonstrating the high heterogeneity of usages for a given identifier resource. Such cross-registry mappings and synonyms in the Bioregistry enable it to act as an interoperability layer to standardize across a large number of external registries and non-registry resources.

### Web portal for interactive and programmatic use

The contents of the Bioregistry can be browsed interactively through the web portal at https://bioregistry.io shown in Figure 2. The portal implements a powerful search feature to help users look up prefixes and CURIEs they encounter in various databases, ontologies, and other biomedical resources. The search feature extends to not only the prefix, synonyms, title, and description of each record, but also all of the corresponding fields in linked records from external registries (Figure 2A). The full prefix list can be browsed (Figure 2B) and each prefix page organizes and contextualizes all information available from novel curation in the Bioregistry as well as imported from external registries (Figure 2C, D). Notably, it links to external registry pages when mappings are available. For example, the page for NCBITaxon (https://bioregistry.io/ncbitaxon) links to a large number of external registry pages (Figure 2E). In addition to the data in the Bioregistry, this page constructs example URIs for first-party providers, third-party providers (e.g., OntoBee^27^, OLS), and external resolvers (e.g., Identifiers.org, N2T) using a combination of information stored about external registries and programmatic logic in the underlying Bioregistry Python package (Figure 2F). The web portal provides several other features including generating pages for each of the external registries integrated into the Bioregistry that show their various properties and functionalities, facilitating curating and displaying user-generated collections of prefixes such as the list of Semantic Web prefixes at https://bioregistry.io/collection/0000002, and listing contributors. The portal also implements a resolver that allows for the uniform construction of URIs from CURIEs that are automatically redirected to the appropriate location based on the URI format string annotated to the CURIE’s prefix. The Bioregistry’s resolver uses the URI scheme https://bioregistry.io/<prefix>:<local-unique-identifier>, similar to the resolver schemes used by Identifiers.org and N2T. Bioregistry also makes available a programming language-agnostic RESTful interface that gives access to all functionality (e.g., search, autocompletion, record retrieval, URI generation) and is documented with OpenAPI/Swagger at https://bioregistry.io/apidocs. The underlying data used to generate each page can be downloaded in a variety of formats including JSON, YAML, TSV, RDF, and others (where applicable). Finally, the portal serves as a hub for links to the code, data, documentation, and narrative surrounding the Bioregistry.

As a companion to the main Bioregistry site, a static site is automatically generated and deployed to https://biopragmatics.github.io/bioregistry using GitHub’s infrastructure. Notably, this includes a *health report* that runs a weekly check for which of the Bioregistry’s resources’ homepages are still accessible (i.e., do not return HTTP 404 or other connection errors) and which resources’ URI format strings are still valid. The site both provides a high-level summary of which resources have recently become invalid as well as a detailed, color-coded table reflecting the statuses of all Bioregistry records. Ultiamtely, this site can help more systematically monitor and improve the maintenance of biomedical resources.

### Exported artifacts for data integration and reusability

The Bioregistry GitHub repository contains the root content of the database (i.e., the registry, metaregistry, and collections) as JSON files in a version controlled setting, serving as the single source of truth. In addition, it makes available several derived artifacts that are meant to facilitate integration with downstream systems and resources. These exported artifacts are regenerated daily, and made available via the web portal at http://bioregistry.io/download and are archived on Zenodo^38^.

In addition to the native JSON format, the Bioregistry data is made available as a set of YAML and TSV files to facilitate reuse. Further, equivalence mappings between resources in external registries are exported into the Simple Standard for Sharing Ontological Mappings (SSSOM)^36^ format. SSSOM is a standard for sharing mappings between different namespaces that we use to represent mappings between resources appearing in different registries, such as the relations exemplified in Figure 1C.

The Bioregistry also provides a number of artifacts to facilitate integration with Semantic Web contexts and linked open data. First, we constructed an RDF schema for the Bioregistry that reuses elements from common Semantic Web vocabularies (e.g., DC, FOAF) and creates its own elements in the bioregistry.schema vocabulary described at https://bioregistry.io/schema. All components of the Bioregistry (i.e., the registry, metaregistry, collections) were jointly exported into RDF under this schema in several commonly used formats including N-Triples, Turtle and JSON-LD. This allows the Bioregistry to be loaded using triple stores (e.g., Virtuoso) or programming libraries (e.g. Python’s RDFLib) and subsequently queried with SPARQL. We also assembled a network derived from the RDF export that can be browsed interactively on the Network Data Exchange (NDEx)^44^ available at https://bioregistry.io/ndex:860647c4-f7c1-11ec-ac45-0ac135e8bacf.

Finally, the Bioregistry makes available several Semantic Web contexts that each map a set of prefixes (e.g., chebi) to a corresponding URI prefix (e.g., http://purl.obolibrary.org/obo/CHEBI_). These are derived from the root data using a set of policies for choosing a preferred prefix and URI format for resources in the registry. In addition to a general purpose context encompassing all of Bioregistry, we make available application-specific semantic contexts for integration with the OBO Foundry, and a context limited to prefixes useful for general Semantic Web resources.

### Maintenance model and governance

In contrast to the maintenance and governance structures employed by existing registries, the Bioregistry takes an alternative approach relying on open data, open code, open infrastructure, automated testing, and automated updating. Similar models have been adopted with great success in existing large collaborative projects such as the OBO Foundry. We accomplished this through several steps. First, the Bioregistry data is stored and versioned using GitHub (see Methods). Second, anyone can propose additions or changes either directly by submitting a pull request to the Bioregistry repository or by filling out an appropriate issue template that triggers an automated generation of a pull request. Both create an open forum for discussion that invites a wide variety of stakeholders to engage (implementation details in Methods). Third, using GitHub allows for the technical implementation of quality control and quality assurance workflows that are coupled to pull requests in order to ensure that all changes meet a predefined set of standards, which are described explicitly and publicly in the contribution guidelines^1^ as well as implicitly in the implementation of the quality assurance workflow. In addition to the open data, open code, open infrastructure philosophy, the Bioregistry project has sought out community guidance on how to establish a governance model that is more robust to the fluctuation of funding and personnel who are actively working on and moderating the project. This has resulted in the establishment of a Review Team and a Development Team as well as a public minimal governance model^2^ that describes how to induct new members, how to remove members, who respectively has the technical authority and community responsibility to facilitate and ultimately judge changes to the underlying database and make changes to the code base, and how to improve the governance model over time. These teams have been initially seeded with members from diverse scientific backgrounds, locations, and institutions to further promote the durability of the project. These guidelines also include a liberal policy on authorship to further demonstrate the project’s commitment to inclusivity.

## 3 Use cases and integrations

Here, we highlight several projects and standards that have already adopted various functionalities of the Bioregistry.

### Supporting Interoperable Data Annotation

Several projects use the Bioregistry to create *prefix maps*, or mappings between prefixes (e.g., uniprot) and their corresponding URI prefixes (e.g., https://www.uniprot.org/uniprot/). These support the the interoperability of data annotations and the conversion between URIs and CURIEs in Semantic Web applications. The Simple Standard for Sharing Ontological Mappings (SSSOM)^36^ is a metadata standard for various mappings (e.g., equivalences) between ontology and database terms and an associated toolset (https://github.com/mapping-commons/sssom-py) based on LinkML (https://linkml.io) for loading, validating, and converting SSSOM content. The standard is meant to encourage higher quality curation in biomedical ontologies which often lack important metadata such as the mapping type, a standardized prefix and local unique identifier for the subject and object terms, provenance about how the mapping was generated, and provenance about who generated the mapping. The default prefix map used in validation is generated by the Bioregistry following the procedure described in Supplementary Section 6.1.

Manubot^34^ is a tool for open collaborative writing that aims to bring automation, customizability, and transparency to scholarly publishing. With Manubot, users write manuscripts using markdown with special support for citation by persistent identifiers represented as CURIEs such as [@doi:10.1371/journal.pcbi.1007128] (which can then be automatically turned into a full citation). Embedding CURIEs in manuscripts is especially valuable when referring to resources that are not citable manuscripts such as clinical trials (e.g., in a review of COVID-19^45^). Manubot initially added support for 700 CURIE prefixes by incorporating Identifiers.org but later switched to the Bioregistry which at the time added support for an additional 365 prefixes. Besides being more comprehensive, the Bioregistry’s open contribution model allowed for addressing several longstanding issues with Identifiers.org including invalid regular expression patterns, missing prefixes, as well as inconsistencies due to some namespaces being redundantly embedded in identifiers.

We discuss the plans and considerations for adopting the Bioregistry for interoperable data annotation in further software and resources including the Biolink Model^15^ and the Alliance of Genome Resources^46^ in Supplementary Section 7.

### Validation and Quality Control of Entity References

Several projects use the Bioregistry to standardize or validate prefixes and local unique identifiers and promote interoperability and reusability. Biomappings^35^ (https://github.com/biopragmatics/biomappings) is a repository for curated and predicted mappings between equivalent (or otherwise related) biomedical entities in different identifier resources. It contains several workflows for generating predicted mappings using Gilda^47^ and provides a web-based curation interface for reviewing predicted mappings and adding novel ones. Biomappings ensures data integrity by validating all prefixes and local unique identifiers in the repository using the Bioregistry. Further, the curation interface uses the Bioregistry to generate links to a web page describing each biomedical entity, making curation easier. Biomappings also generates a web-based summary of its content that uses the Bioregistry to provide links to identifier resources and to resolve CURIEs.

The Integrated Network and Dynamical Reasoning Assembler (INDRA)^37^ assembles biomedical knowledge from multiple databases combined with text mining of scientific publications to construct executable models. When performing assembly, INDRA maintains references to biomedical entities that are grounded to one or more identifier resources. It uses the Bioregistry to first check that the prefixes used in these groundings are standardized, and then to validate the associated unique local identifier according to the pattern provided by the Bioregistry. This validation is critical for maintaining consistency in INDRA’s automated assembly workflows.

The Phenotype Knowledge Translator (PheKnowLator)^33^ ecosystem constructs FAIR biomedical knowledge graphs using ontologies and reasoning with the addition of non-ontological data sources. PheKnowLator uses the Bioregistry to standardize references in CURIEs and URIs from both data types to provide semantically consistent results for downstream use cases. The Bioregistry helps overcome significant challenges posed by ontologies (e.g., changing namespaces over time, data that is not from an ontology that does not provide valid namespaces or URIs), and their integration. The Bioregistry API has become a vital component of the build process and is used to standardize URIs for all entities and triples. It has also provided new opportunities to extend PheKnowLator’s testing harness. Overall, the inclusion of Bioregistry has improved the PheKnowLator Ecosystem and the knowledge graphs it produces. Similarly, the Graph Representation leArning, Predictions and Evaluation (GRAPE)^48^ software package uses the Bioregistry to normalize the identifiers in several networks and knowledge graphs (including PheKnowLator).

### Contextualizing Entities with Website Links

Several projects use the Bioregistry to generate and resolve URIs within their APIs or user-facing websites in order to provide additional context to the entities they reference. BridgeDb^32^ is a web service that maps between local unique identifiers from different identifier resources representing equivalent entities (e.g., P0DP23 in UniProt Q17855525 and in Wikidata for the Calmodulin-1 protein). The Bioregistry has been integrated in BridgeDb’s Java and R clients as well as Bacting^49^ to enable lookup based on standardized CURIEs, to enable creating internal BridgeDb identifier objects *via* standardized CURIEs, and to generate URIs resolvable through the Bioregistry web application.

The Bioregistry has also been used in several websites to generate URLs for human genes, protein complexes, and other entities. For example, the DUB Portal^50^ is a website summarizing experimental analyses of deubiquitinating enzymes and uses the Bioregistry to link to human genes in HGNC and protein families in the FamPlex vocabulary^51^. The website for FamPlex^3^ also uses the Bioregistry to standardize and link references for human genes; equivalence mappings to InterPro^52^, Medical Subject Headings (MeSH)^53^, GO, Complex Portal^54^, NextProt^55^ for protein families and complexes; and references for publications in PubMed and PubMed Central. Similarly, the interactive user interface for the BERN2^56^ named entity recognition platform standardizes its biomedical entities and generates links using the Bioregistry.

### Unified Access to External Registries

Because of its integration of external registries, the Bioregistry is also useful for unified access to their respective data. The OBO Foundry^25^ facilitates the coordinated development of biomedical ontologies through a set of guiding principles and community organization. Its associated repository (https://github.com/OBOFoundry/OBOFoundry.github.io) stores the structured metadata about each ontology, including their preferred prefix, title, homepage, description, and usages. The Bioregistry is used to support the standardization and maintenance of this metadata in several ways described in detail in Supplementary Section 6.2.

## 4 Discussion

We presented the Bioregistry, an integrative registry of biomedical identifier resources. The Bioregistry takes a novel approach to curation by importing and harmonizing data from external registries that can be further improved and extended with novel curation. It relies on an open data, open code, open infrastructure philosophy combined with a novel governance strategy to foster community contributions and engagement, and ensure its longevity and adoption. It uses public infrastructure for quality assurance, distribution, and deployment to promote transparency, reduce cost, and uncouple its long-term maintenance from a specific institution, funding source, or group of maintainers. While the Bioregistry demonstrates higher coverage and metadata granularity than other registries, it also explicitly encourages reuse and redistribution via its highly permissive CC0 license.

### Limitations

Entries in the Bioregistry that represent identifier resources, their preferred CURIE prefix, and other metadata are integrated semi-automatically from external registries (such as Identifiers.org and Name-to-Thing (N2T)^28^) or manually curated directly in the Bioregistry. The design choice that the Bioregistry semi-automatically imports and aligns content from external registries is important for maintaining broad coverage, and to distribute curation effort across multiple projects. However, this still poses challenges for consistency. Namely, the Bioregistry has limited ability to enforce guidelines and conventions in other registries. For instance, there are differing views in the community on stylistic choices in the capitalization of preferred CURIE prefixes (e.g., chebi vs CHEBI or ChEBI) for identifier resources. Drawing on other registries can also lead to future conflicts where multiple registries choose the same CURIE prefix for two different identifier resources, creating a situation that has to be retroactively arbitrated in the Bioregistry (an example is given in Supplementary Section 5.2). Nevertheless, the Bioregistry maintains guidelines^4^ for creating new identifier resource prefixes, which, if followed, can mitigate these issues. Further, the purview of the Bioregistry does not extend to directly advising and mentoring creators of new identifier resources to make good choices in their identifier schemes. Creators of such resources can rely on recommendations such as those suggested by McMurry *et al*. (2017)^6^.

Adopting the Bioregistry’s standard for prefixes, CURIEs, and URIs in a new resource is straightforward. However, applying it retroactively to an existing resource can pose challenges. It may require updating the data and associated code in the resource itself as well as in downstream consumers of the resource. This can manifest in several ways, including updating non-standard synonyms (e.g. many ontologies use MSH as a non-standard prefix for MeSH), updating non-standard construction of CURIEs (e.g., using redundant prefixes as prescribed by Identifiers.org like in GO:GO:0006915), or updating non-standard URIs (e.g., switching all ORCID URIs to use the *https* protocol). If such changes are not feasible in the resource, it is still possible to implement mappings to the Bioregistry or create custom exports following the Bioregistry standard, potentially broadening the resource’s interoperability.

The Bioregistry provides a solution for standardizing references to individual entries in identifiers resources. However, it is often the case that multiple identifiers resources contain entries representing equivalent entities (e.g., multiple disease ontologies representing the same disease) leading to redundancy when integrating disparate resources, such as when constructing knowledge graphs. Determining which identifier resource to prioritize when representing an entity that appears in multiple resources is beyond the scope of the Bioregistry. Nevertheless, the Bioregistry can contribute to the standardization of the cross-references between equivalent entities in different identifiers resources (cross-references, in practice, often use non-standard CURIEs and URIs) thereby helping redundancy resolution among them.

While the Bioregistry is limited to resources of interest to researchers in the life sciences, its methodology and technological implementation could extend to other scientific areas. Ultimately, the Bioregistry could serve as a template for the creation of domain-specific metaregistries in other areas or be the basis for the creation of a metaregistry spanning multiple scientific domains.

### Future Work on the Bioregistry

Following the initial development, deployment, and early adoption of the Bioregistry, two ongoing challenges remain. The first is to be responsive in the maintenance, enrichment, and extension of the content in the registry to best reflect the reality of the ever-changing landscape of biomedical identifier resources. While this has not been realized by previous registries, the Bioregistry’s combination of technical infrastructure and governance model will enable this effort in a sustainable way. Further, we plan to develop tools and curation practices to proactively, systematically identify new resources to add to the Bioregistry.

The second is to build and maintain a community of users. This entails continuing to engage multiple groups of users and stakeholders. This includes curators, and consumers of biomedical resources, as well as groups designing automated data- or knowledge-extraction and aggregation systems. Serving the needs of these communities requires identifying their challenges, and improving the Bioregistry’s data model, tooling, and content accordingly. It also entails facilitating discussion between a diverse set of individuals and offering training for usage of the Bioregistry and its philosophy. To this end, the authors plan to organize a set of recurring community workshops (following an initial workshop held in 2021^5^) around the topics of identifier resources and registries.

### Future Vision

We envision the Bioregistry could more broadly be used to promote and support the standardized annotation of data, models, ontologies, and scientific literature. First, the growing body of data being made available through publications and data repositories often lack standardized annotations to their records (e.g., columns in a table, nodes/edges in knowledge graphs). If adopted by the diverse set of curators, developers, maintainers, and users of life science tools and resources, the Bioregistry could provide a consistent way of annotating these data to make them more FAIR, especially facilitating reuse.

Second, we envision the Bioregistry supporting the standardization of structured metadata associated with models and networks derived from data such as mechanistic models (e.g., in the BioModels database^57^), network-based models (e.g., in Network Data Exchange (NDEx)^44^), knowledge graphs (e.g., those described by Bonner *et al.^58^)*, and machine learning models (e.g., such as those trained by GRAPE^48^) in order to promote their interoperability and reuse. For example, despite the recent proliferation of biomedical knowledge graphs^58^, there has been little convergence on standardized syntax or semantics for identifying nodes and edges. The Bioregistry can support this effort both on a low level and also by integrating into higher-level tools that generate and exchange graph-like data such as KGX and Biolink (see Supplementary Section 7.2) that support a larger variety of use cases. More generally, we believe that this will enable the growing audience of machine learning researchers who are interested in combining biomedical datasets in novel ways.

This vision aligns well with the recommendations from a recent assessment of the reproduciblity of such models^59^ that highlighted the more general importance of annotation using high-quality controlled vocabularies like GO and ChEBI.

Third, though biomedical ontologies have proven invaluable for data annotation, key ontologies still suffer from a lack of standardization of cross-references^25^, making it difficult to merge and reason across ontologies and other structured data sources. Given that ontologies are often curated in public version-controlled repositories in standardized formats (e.g., OBO, OWL), the Bioregistry could be used to support their semi-automated standardization and maintenance in order to both reduce curation burden and potentiate their value in data integration scenarios.

Finally, we envision the potential adoption of the Bioregistry by academic publishers to support the standardized annotation of named entities in the text provided by authors (e.g., the BioFactoid^60^), and thereby decrease the need for doing expensive and error-prone post-processing like automated named entity recognition on publications to create structured representations.

## Methods

The Bioregistry repository tightly couples the data to a Python package that facilitates loading, accessing, and modifying the root data files. It provides several high-level data structures and workflows for accessing and reasoning over the Bioregistry and external registries’ integrated data, that support the quality assurance workflows (described above), the web application (described above), the alignment workflows (described below), the generation of derived artifacts (described above) and other user-facing functionality such as prefix standardization, CURIE standardization, and URI parsing. Full documentation for the Python software package can be found at https://bioregistry.readthedocs.io.

### Alignment

While manual curation of mappings to external registries is feasible when adding novel prefixes to the Bioregistry, the frequency of updates to external registries motivated the development and application of periodic automated and semi-automated alignment.

We first stratify all external registries into three categories based on their available metadata, biomedical scope, focus on assigning global prefixes to resources, and governance. The first group with metadata availability, a biomedical scope, and focus on assigning global prefixes contains the Identifiers.org, the OBO Foundry, OLS, N2T. The second group contains registries such as GO, NCBI, UniProt, Cellosaurus, and FAIRSharing that contain entries that do not correspond to identifier spaces which are excluded from the import. It additionally included registries like BioContext and BioPortal because of insufficient metadata that often made it impossible to determine what identifier resource the metadata refers to. The third group contains registries with minimal metadata or lack of biomedical focus such as Prefix.cc. The alignment algorithm first generates a lookup table based on the canonical prefix, preferred prefix, and all prefix synonyms (see Supplementary Section 3 for details on the data model and Supplementary Figure 2 for a schematic diagram of this process) for each resource in the Bioregistry. The prefix policies and automated quality assurance checks in the Bioregistry ensure that there are no collisions in this lookup table. For each external registry, the data are downloaded, normalized, and exactly one field is annotated as the external prefix (see Supplementary Figure 1). All Bioregistry prefixes that already have been mapped to an external prefix in the external registry are removed from the lookup table. Similarly, all external prefixes that already have mappings are not considered for new mappings. Each external prefix that matches an entry in the lookup table is assigned an automated mapping. A manually curated list of incorrect mappings and collisions are used to post-process the automated mappings and remove false negative mappings (see Supplementary Section 5.2). External prefixes that could not be mapped to a Bioregistry prefix are handled based on their stratification. For the first group of registries, the prefix is added as a new record to the Bioregistry. For the second group of registries, the prefix is added to a curation sheet along with its relevant metadata (e.g., title, homepage, example identifier) for later manual curation. For the third group of registries with minimal metadata, no report is made.

### Promoting sustainability and longevity through automation

The Bioregistry is hosted on GitHub (https://github.com/biopragmatics/bioregistry) to take advantage of its public, cloud-based version control, collaboration, and workflow management platforms. The single source of truth data (i.e., root data) for the Bioregistry is stored in version control. This implicitly versions all minor changes with git commit hashes and allows git tags to be used to mediate releases, which are automatically archived and re-distributed on both GitHub and Zenodo (https://zenodo.org).

The Bioregistry uses GitHub Actions as a continuous integration service to run code and data quality assurance to promote the maintainability and integrity of the resource (see Supplementary Section 4.1). They further enable workflows for automatically generating pull requests and notifying reviewers to enable non-technical users to make submissions to the resource. The Bioregistry further uses GitHub Actions as a continuous delivery and continuous deployment system to run the aforementioned alignment workflows, generate derived artifacts, release code to the Python Package Index (PyPI), containerize code on Docker Hub (https://hub.docker.com), deploy the web application to Amazon Web Services (https://aws.amazon.com) on a daily basis (see Supplementary Section 4.2). Combined, the continuous integration, delivery, and deployment services allow contributors and consumers of the Bioregistry to more easily propose improvements, review them as a community, and see them reflected in the data and website without the need for manual intervention by the project team. Using an entirely free, public, and open public infrastructure to do so promotes longevity and sustainability by mitigating the monetary requirements. Further, the technical requirements of the deployment of the web service and hosting are also minimized such that hosting costs around 33$/year and compute costs around 27$/year (see Supplementary Section 4.3).

## Code availability

The source code for the Bioregistry is available at https://github.com/biopragmatics/bioregistry under the MIT License. The source code specific to the version of Bioregistry used in this article (v0.5.132) is archived on Zenodo^38^.

## Data availability

All data analyzed during this study are available on GitHub at https://github.com/biopragmatics/bioregistry. The data specific to the version of Bioregistry used in this article (v0.5.132) is archived on Zenodo^38^.

## Acknowledgements

CTH, KK, BMG were funded under the Defense Advanced Research Projects Agency (DARPA) Young Faculty Award [W911NF-20-1-0255]. EW received funding by NWO grant [203.001.121].

The authors would like to acknowledge the developers, maintainers, and curators of each of the external registries referenced throughout this manuscript and used in the Bioregistry, whose prior work enabled ours. More detailed “live” acknowledgments for these registries can be found at https://bioregistry.io/acknowledgements.

## Author contributions statement

CTH and BMG conceived and implemented the software and resource, analyzed the results, and wrote the manuscript. CJM, JK, JM, and MAH contributed to the conception and design of the resource and software. AR, BMG, CTH, CJM, DDF, DSH, DW, MB, NM, SM, TL, and TJC performed data curation for the resource, BMG, CTH, DRU, EW, HBH, KK, NM, and SM contributed to the software. CJM, DSH, DRU, EW, JM, NM, SM, TJC co-wrote the manuscript. All authors reviewed and edited the manuscript.

## Footnotes

1 https://github.com/biopragmatics/bioregistry/blob/main/docs/CONTRIBUTING.md

2 https://github.com/biopragmatics/bioregistry/blob/main/docs/GOVERNANCE.md

3 https://sorgerlab.github.io/famplex

4 https://github.com/biopragmatics/bioregistry/blob/main/docs/CONTRIBUTING.md#submitting-new-prefixes

5 https://biopragmatics.github.io/workshops/WPCI2021.html

