## Supplementary Information for "Unifying the Identification of Biomedical Entities with the Bioregistry"

### Supplemental Information for The Bioregistry

#### 1 URIs and CURIEs have high heterogeneity

The introduction of the main text alludes to the high number and heterogeneity in uniform resource identifiers (URIs) and compact uniform resource identifiers (CURIEs). Table 1 and Table 2 demonstrate this phenomena using the `P0DP23` entry in the UniProt Knowledgebase representing the Calmodulin 1 as an example.

| URI | Description | Type | Use Case(s) |
|---|---|---|---|
| http://purl.uniprot.org/uniprot/P0DP23 | UniProt persistent URL | First-party | Linked open data |
| https://www.uniprot.org/uniprot/P0DP23 | UniProt protein page | First-party | User-friendly navigation |
| https://identifiers.org/uniprot:P0DP23 | Identifiers.org URI using HTTPS | Third-party | CURIE resolution, |
| http://identifiers.org/uniprot:P0DP23 | Identifiers.org URI using HTTP | Third-party | ... mathematical modeling, |
| https://identifiers.org/uniprot/P0DP23 | Legacy URI using HTTPS | Third-party | ... biocuration, |
| http://identifiers.org/uniprot/P0DP23 | Legacy URI using HTTP | Third-party | " |
| https://n2t.net/uniprot:P0DP23 | Name-to-Thing URI | Third-party | CURIE resolution |

**Table 1.** Example URIs for the Calmodulin 1 protein (P0DP23) in the UniProt Knowledgebase.

| CURIE | Prescribed by Registries |
|---|---|
| `UniProtKB:P0DP23` | Cellosaurus, Gene Ontology, Prefix Commons |
| `uniprot:P0DP23` | Identifiers.org, N2T |
| `UniProt:P0DP23` | NCBI GenBank |

**Table 2.** Example CURIEs for the Calmodulin 1 protein (P0DP23) in the UniProt Knowledgebase. Figure 1C in the main text demonstrates an even more heterogeneous scenario for CURIEs for entries in the NCBI Taxonomy Database.

#### 2 Comparsion of External Registries

In addition to comparison of the data models of external registries in the main text, this section compares the governance and maintenance models of each external registry in Table 3. Below are detailed descriptions of fields appearing in the table.

**Scope** This field denotes the scope of prefixes which the registry covers. For example, some registries are limited to ontologies, some have a full scope over the life sciences, and some are general purpose.

**Status** This field denotes the maintenance status of the repository. An active repository is still being maintained and also is responsive to external requests for improvement. An unresponsive repository is still being maintained in some capacity but is not responsive to external requests for improvement. An inactive repository is no longer being proactively maintained (though may receive occasional patches).

**Imports External Prefixes** The imports external prefixes column denotes if the registry reuses and harmonizes content from external registries. For example, Name-to-Thing (N2T)[14] and the Bioregistry import prefixes and their metadata. Similarly, lookup services like Ontobee, Ontology Lookup Service (OLS)[17], and AberOWL import the OBO Foundry ontologies in bulk.

**Accepts External Contributions** This field denotes if the registry (in theory) accepts external contributions, either via suggestion or proactive improvement. This field does not pass judgement on the difficult of this process from the perspective of the submitter nor the responsiveness of the registry. This field does not consider the ability for insiders (i.e., people with private relationships to the maintainers) to affect change.

**Public Version Control** This field denotes if the registry stores its data/code in publicly available version control system, such as GitHub or GitLab. Currently there is no resource that does one but not the other, so this is grouped (for now).

**Issue Tracker** This field denotes the public issue tracker for issues related to the code and data of the repository.

| Registry | Scope | Status | Imports External Prefixes | Curates Novel Prefixes | Accepts External Contributions | Data Public Version Control | Public Issue Tracker | Has Public Review Team |
|---|---|---|---|---|---|---|---|---|
| Bioregistry | Life Sciences | Active | Y | Y | Y | Y | Y | Y |
| AberOWL[1] | Bio-Ontologies | Active | Y | | | | | |
| AgroPortal[2] | Agro-Ontologies | Active | | Y | Y | | | |
| BioContext[3] | Internal | Active | | Y | Y | Y | Y | Y* |
| Biolink[4] | Internal | Active | | Y | Y | Y | Y | Y* |
| BioPortal[5] | Bio-Ontologies | Active | | Y | Y | | Y | |
| Cellosaurus[6] | Internal | Active | | Y | | | | |
| CHEMINF[7] | Chemistry | Unresponsive | | Y | Y | Y | Y | Y* |
| CropOCT[8] | Agro-Ontologies | Active | | Y | Y | | Y | Y |
| EcoPortal[9] | Eco-Ontologies | Active | | Y | Y | | | |
| FAIRSharing[10] | Life Sciences | Active | | Y | Y | | | |
| GO[11] | Internal | Inactive | | Y | Y | Y | Y | Y* |
| HL7/FHIR[12] | General | Active | | Y | Y | | | |
| Identifiers.org[13] | Life Sciences | Unresponsive | | Y | Y | | Y | |
| N2T[14] | Life Sciences | Active | Y | | | | | |
| NCBI GenBank[15] | Internal | Inactive | | Y | | | | |
| OBO Foundry[16] | Bio-Ontologies | Active | | Y | Y | Y | Y | Y |
| OLS[17] | Bio-Ontologies | Active | Y | | Y | Y | Y | Y* |
| OntoBee[18] | Bio-Ontologies | Active | Y | | Y | | Y | |
| Prefix Commons[19] | Life Sciences | Active | | Y | Y | | | Y |
| UniProt[20] | Internal | Active | | Y | | | | |
| Wikidata[21] | General | Active | | Y | Y | | | Y* |

**Table 3.** A survey of registries' governance and maintenance models. A detailed description of each column can be found in Supplementary section 2. Asterisks in the "has public review team" column denote that rather than an explicit list, the reviewers can be inferred through the registry's version control system.

**Public Review Team** This field denotes if the registry's reviewers/moderators for external contributions known. If there's a well-defined, maintained listing, then it can be marked as public. If it can be inferred, e.g. from reading the commit history on a version control system, then it can be marked as inferrable. A closed review team, e.g., like for Identifiers.org can be marked as private. Resources that do not accept external contributions can be marked with N/A. An unmoderated registry like Prefix.cc is marked with 'democratic'.

#### 3 Data Model

This section describes the data model in detail. This may be subject to change following publication, and will be maintained as a living document at `https://github.com/biopragmatics/bioregistry/blob/main/docs/datamodel.md`.

##### 3.1 Metadata and Properties

**Prefixes** Each entry in the Bioregistry is annotated with a required canonical prefix (i.e., lower case, containing no strange punctuation or characters), an optional preferred prefix (i.e., containing stylization), and an optional list of alternate prefixes (i.e., synonyms).

**Status** Each includes flags if the resource for the prefix is deprecated (e.g., aero), proprietary (e.g., namerxn), or doesn't contain any terms (e.g., chiro). These flags are helpful in subjectively deciding which resources could be considered as reliable.

**Name, Description, and Homepage** All entries in the Bioregistry must have a name and description. This is important not only for active resources, but also for deprecated resources so the integrative registry can be used as a research tool and reference such as when encountering the variety of historically obscured cross references in ontologies in the Open Biomedical Ontologies (OBO) Foundry[16]. All active resources must additionally have a homepage, while inactive resources may not have a homepage due to their respective sites being taken down. Entries are highly encouraged to include reference URLs and contributor free text comments, especially for deprecated resources to give context to readers.

**License** License information is only readily available via the OBO Foundry and the OLS. Most use permissive licenses from the Creative Commons (either CC-BY or CC-0). Some licenses include license versions and some do not. There are several instances of conflict, often due to specification of the version of the Creative Commons license. Licenses only appearing a small number of times, such as CC-BY-SA, CC-BY-NC, and CC-BY-NC-SA were collapsed into "Other". Licenses that were not appropriate for data (e.g., variants of the Apache License, GNU GPL) and custom licenses (e.g., in the case of the Human Phenotype Ontology) were also collapsed into "Other".

**Version** The OLS is the only registry that consumes the data it references and provides detailed accessible artifacts, and is therefore the only registry that reports version information. Other lookup services like Aber-OWL[1] and OntoBee[18] consume ontologies but do not generate metadata reports. The OBO Foundry also references versioned data, but does not consume it and therefore can not report version information. Wikidata also contains version information for some databases, but is not currently viable for generally tracking version information. The other registries (e.g., Identifiers.org, N2T) do not report version information as their resolution services are independent of the data versions. Alternatively, the Bioversions project sets out to be a registry-independent solution for identifying current versions of different databases, ontologies, and resources.

**Local unique identifiers (LUIDs)** All non-deprecated entries in the Bioregistry must include one or more example LUIDs. While each optionally (highly recommended) includes a regular expression pattern describing local unique identifiers, there are cases when they are difficult to generate due to the high complexity and heterogeneity of identifiers (e.g., Ensembl identifiers are highly complex), the lack of enough examples as is often the case with deprecated identifiers, or the triviality of assigning a wildcard pattern to a small enumerated namespace like FOAF, RDF, or RDFS.

While the Bioregistry imports identifier regular expression patterns from several registries (i.e., Identifiers.org[13], N2T, Prefix Commons[19], the Gene Ontology (GO)[11], and Wikidata), there exist many philosophical and practical discrepancies. A major one arises in the definition and management of redundant prefixes embedded in identifiers that is common to ontologies. For example, the colloquial local unique identifier for apoptosis in GO is `GO:0006915`, so the corresponding prefixed CURIE for this entity is `GO:GO:0006915`. Each registry handles cases like this slightly differently, whether it is to include the redundant prefix capitalized in the pattern, to include an extra flag in the metadata associated with the prefix, or whether to completely handle this programmatically.

In order to promote backwards compatibility with some Identifiers.org prefixes, entries can contain an annotation "namespace embedded in LUI" to allow for overriding of Identifiers.org metadata as well as the "banana" which explicitly states how the namespace embedded in LUI appears, for situations where it is not the same as the prefix itself (e.g., `HOG`).

Because there is such low consistency, the Bioregistry introduces a new field called the *banana* where the potentially redundant prefix can be explicitly enumerated, which allows for a general solution that can apply to `FBbt`, `VariO`, and other mixed-case embedded prefixes. The Bioregistry Python package includes functions for reformatting CURIEs based on the desired context (e.g., for general use, for compatibility with Identifiers.org). Further discussion on this topic can be found at `https://github.com/biopragmatics/bioregistry/issues/191`.

**Provider (URI Format String)** Each entry optionally (highly recommended) includes one or more URI format strings that can be used to generate URIs for a local unique identifier. All active entries should have a provider, if possible. Deprecated resources likely do not have providers. Two persisting issues are due to providers being decommissioned but not removed from registries and the propagation of resolver services as providers which exacerbates the first. For example, because many ontologies use OBO-like persistent uniform resource locators (PURLs), they are often annotated in registries as providers even though they do not resolve to anything. It is a future goal to provide more "health" checks over each registry and the Bioregistry as a whole.

**Availability** Each entry includes three optional fields for when the resource is available as an ontology in the OWL (`https://www.w3.org/TR/owl2-syntax`), OBO (`https://owlcollab.github.io/oboformat/doc/obo-syntax.html`), or OBO Graph JSON (`https://github.com/geneontology/obographs`) formats. These entries are typically imported from the OBO Foundry and OLS and are manually annotated to support large-scale ontology acquisition and processing such as with ROBOT, Pronto, or PyOBO.

**Attribution** Each includes two required attribution fields for the contributor and reviewer that each require a minimum of an ORCiD identifier and name with optional email address and GitHub handle. Each also includes an optional attribution field for an external contact person for the resource that could have either the ORCiD identifier, email, GitHub handle.

###### *3.1.1 Ontological Relationships Between Prefixes*

In addition to metadata and properties, the Bioregistry includes rich ontological relationships between prefixes in the Bioregistry as well as external prefixes.

**Exact Matches** The most novel aspect of the Bioregistry is its ability to store equivalence mappings (e.g., `skos:exactMatch`) between Bioregistry records and external records (external records' semantics are mediated by the metaregistry). Each entry in the Bioregistry can contain several mappings to different databases. Typically, each prefix can only have one exact match in each database. Exceptions have arisen due to duplicate records, in which case the mapping is curated to the canonical external record. These records support interoperability by enabling conversion between the standard flavor of a prefix defined by the Bioregistry and context-specific variants.

**Depends On** Each entry contains a list of external entries that its associated resource depends on. This is particularly useful in ontologies, since they may either import terms from an external ontology or use external prefixes in their xrefs, provenance, or relationships. These are mostly imported from the OBO Foundry but also have an aspect of novel manual curation. While not explicitly stored in the source data, the Bioregistry python package infers the inverse relationship ( i.e. appears in) for easy access given a prefix.

**Provides** While prefixes in the Bioregistry are supposed to correspond to nomenclature authorities, this is not always true because it imports from external sources that don't enforce this constraint. For example, the Comparative Toxicogenomics Database[22] uses NCBI Gene for naming genes and Medical Subject Headings (MeSH)[23] for naming diseases and chemicals. Identifiers.org has minted 3 prefixes (ctd.gene, ctd.disease, and ctd.chemical) that mostly reflect the entries of the authorities for which they are providers. Another example is ValidatorDB[24], which provides information based on Protein Databank[25] records. An even more exotic example are the Gene Ontology Annotations provided by the EBI because it provides for several types of identifiers including those from UniProt[20], RNA Central[26], and the Complex Portal[27].

Therefore, prefixes in the Bioregistry can be annotated with the prefix for which they provide (e.g., ctd.gene provides mesh). Along with the part of and has canonical relationships (described below), this relationship can promote better standardization and help deconvolute multiple prefixes that use the same URI format string, which is problematic when generating high quality prefix maps for use in CURIE-URI interconversion.

**Has Canonical** While there should not be redundancies in the Bioregistry, there are several scenarios in which two or more prefixes equally correspond to the same nomenclature authority. Because these can not be merged without making the data model for the Bioregistry much more complicated and inaccessible, the has canonical annotation allows for the subjective choice of which is considered highest priority. A few scenarios in which this annotation is used are:

1. A prefix has been replaced by another one (e.g., hgnc.genefamily was replaced by hgnc.genegroup)

2. A prefix is redundant of another (e.g., glycomedb is redundant of glytoucan, pdb-ccd is redundant of pdb.ligand)

3. Multiple prefixes are used by different groups for the same shared semantic space, but none of them own it (e.g., insdc.run, ena.embl)

Records in the has canonical relationship do not necessarily have the same URI format string, but if they do, this relationship further promotes choosing a deterministic prefix when parsing an URI in combination with the provides and part of relationships.

**Part Of** There are several flavors of hierarchical relationships between prefixes in the Bioregistry annotated with the part of relationship. For example, chembl.compound and chembl.target are each a part of chembl and kegg.pathway and kegg.ligand are each a part of kegg. Connecting these prefixes provides significantly more context to readers of the Bioregistry. Other scenarios include:

1. no shared prefix, has parent prefix (e.g., fbbt, fbcv, fbrf, ... flybase)
2. has shared prefix, has dot delimiter, has a parent prefix (e.g., kegg with kegg.pathway, kegg.ligand, ...),
3. has shared prefix, has dot delimiter, no parent prefix (e.g., insdc.cds/insdc.gca/insdc.sra),
4. has shared prefix, no dot delimiter, has a parent prefix (N/A)
5. has shared prefix, no dot delimiter, no parent prefix (e.g., dlxb/dlxb, NCBIGene/NCBIProtein/NCBITaxon)
6. prefix matching resource name and extra prefixes (e.g., biogrid and biogrid.interaction)

In several cases such as KEGG and ChEMBL, the parent prefix and child prefixes share a URI format string. Practically, the parent prefix would be sufficient, but it is often pertinent to use a subspace to denote entity types within the nomenclature. In the case of KEGG, each different entity type has a different identifier pattern. In the case of ChEMBL, all different entity types have the same identifier pattern. Ultimately, the part of relationship is the last part combined with the provides and has canonical relationships along with a small amount of additional logic to construct a high-quality prefix map.

The Rat Genome Database[28] (RGD) constitutes an edge case with its three prefixes: rgd, rgd.qtl, and rgd.strain. The rgd prefix is more of a bucket than a parent - it includes all of the entity types (e.g., genes, articles) in the RGD that are neither quantitative trait loci (QTLs) nor strains. Because of cases like this, we have begun discussions on imposing a prefix subspacing policy described at `https://github.com/biopragmatics/bioregistry/issues/133`.

#### 4 Software and Hardware

##### 4.1 Testing Suite

The Bioregistry GitHub repository uses pyroma and check-manifest to check package metadata integrity, black and isort to enforce a consistent code style, flake8 for code quality assurance, mypy for optional static type checking, doc8 and docstr-coverage to ensure properly formatted documentation, sphinx to build inline documentation into external narrative documentation, ReadTheDocs to host documentation, pytest and xdoctests to run unit tests, and codecov to assess testing coverage.

The unit tests not only assess that the code for the Bioregistry Python package and web application work, but more importantly, constitute a technical implementation of the data quality guidelines, which significantly reduces curation burden. For example, when a new prefix is requested via a pull request, the existence of required fields is checked via a unit test and an error report is given to the contributor before needing to request a review. More detailed checks also assert that the homepage does not return HTTP 404, that the given example matches the given regular expression pattern, that the prefix does not have any illegal characters, that acronyms are written out in full in the label, and more.

##### 4.2 Automated Deployment

The Bioregistry code on Python Package Index (PyPI) (`https://pypi.org/project/bioregistry/`) is used to build a Docker container based on the Python 3.10 alpine base image, which significantly reduces non-essential components. The compressed image weighs less than 40 MB of disk space, runs inside Docker with about 65 MB of memory at baseline, and can easily fit on a dedicated t4g.nano instance on Amazon Web Services (AWS) (i.e., the smallest possible) that costs about $37/year on-demand or around $20/year reserved. The image is finally pushed to a public repository on Docker Hub (biopragmatics/bioregistry).

##### 4.3 Hosting

The Bioregistry web application is redeployed on each of two AWS Elastic Cloud Compute (EC2) instances by a script triggered daily that shuts down the existing service, pulls the newly minted image from Docker Hub, and deploys the new service. The application is served via a load balancing service to stay secure and highly available. It is forwarded to the bioregistry.io domain whose registration costs about $33 per year. The SSL/TLS certificate for bioregistry.io (so it can be served with HTTPS) is managed through the AWS Certificate Manager.

#### 5 Alignment

We provide a schematic diagram of the four typical preprocessing steps for external registry metadata in Figure 1:

1. In the example in Figure 1, the data are downloaded in bulk from AberOWL's API endpoint[1]. Other external registries are downloaded in bulk via flat files, by pagination through API endpoints, or via web scraping.
2. The results from AberOWL's API are returned as JSON and can be readily parsed. Other external registries return results in various formats, some which need custom parsers.
3. Custom Python code[2] is used to process each record. In this example, the code renames several fields, applies string normalization to their values, and introspects on the file extensions for the ontology download link.
4. The processed records, which now only contain content relevant for the Bioregistry, are stored under version control and later used in the alignment pipeline outlined in Figure 2.

We provide a schematic of the alignment workflow in Figure 2. While most records have a single prefix that is typically mappable, records like those from FAIRsharing can have synonyms that are useful in alignment. Records in Wikidata are typically not mappable since they use opaque identifiers instead of prefixes.

##### 5.1 Alignment Quantfication

We calculated an upper bound on the number of one-to-one mappings between prefixes in each pair of external registries of 58,058. This decreases by 4.7 times to 12,224 mappings when using the Bioregistry as a mapping hub via its metaregistry. Of these, 5,243 (43%) have been curated and 6,981 (57%) remain, but these numbers are subject to change depending on both updates to the Bioregistry and external registries.

##### 5.2 Alignment Case Studies

For example, the Gene Expression Omnibus has historically used the `geo` prefix and was included in Identifiers.org, Prefix Commons, and ultimately the Bioregistry. When the Geographical Entity Ontology was later registered on the OBO Foundry, the alignment algorithm automatically created a mapping between the geo prefix in the Bioregistry, corresponding to the Gene Expression Omnibus. While the technical solution was to curate the negative mapping in the mismatch list, the community solution required discussion with the Geographical Entity Ontology curators to pick an alternate prefix. Ultimately, it was decided that geogeo could be used for the Geographical Entity Ontology in the Bioregistry, while having an explicit mapping to the geo record in the OBO Foundry. This anecdote demonstrates that the goal of the Bioregistry is not to dictate which prefixes each community uses, but rather provide the infrastructure for mapping between different contexts. As a follow-up, the OBO Foundry instituted a requirement that new ontologies' prefixes do not conflict with existing Bioregistry and Bioportal prefixes.

A related example is CiteXplore, a precursor of the EuropePMC project that often uses ctx as a prefix synonym. This synonym conflicts with the Cerebrotendinous Xanthomatosis Ontology[29] (`https://bioportal.bioontology.org/ontologies/CTX`) in BioPortal, which also uses `CTX`. However, the Cerebrotendinous Xanthomatosis Ontology does not have an entry in the Bioregistry because it is not represented in a preferred registry (described earlier) nor has there been community interest to manually curate an entry on it. The collision for CiteXplore to CTX has been curated to remove this false positive mapping. Overall, collisions occur relatively infrequently in high quality resources, with the false positive mapping file currently only holding a handful of entries.

#### 6 Current Use Cases

This section gives additional details on the use cases presented in the main text.

---

[1] `http://aber-owl.net/api/ontology/?drf_fromat=json&format=json`

[2] `https://github.com/biopragmatics/bioregistry/tree/1e2255291e2b0bd85efff7f152cc83523734351d/src/bioregistry/external`

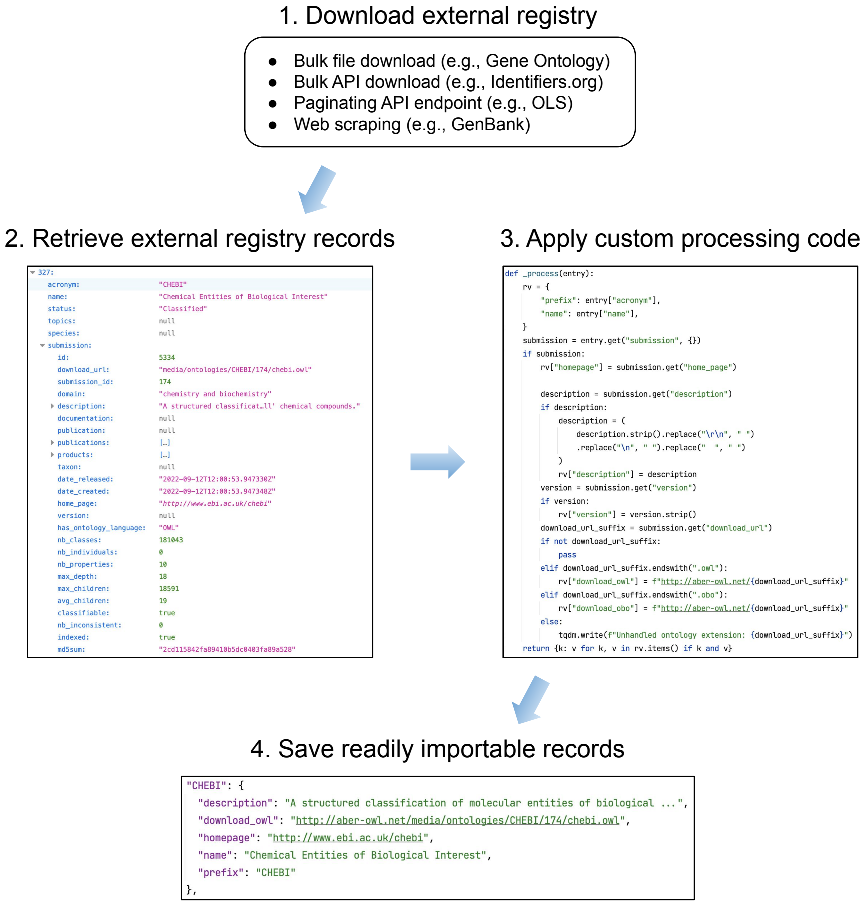


**Figure 1.** A schematic diagram of the external registry download and processing workflow.

##### 6.1 Simple Standard for Sharing Ontological Mappings[30]

While each Simple Standard for Sharing Ontological Mappings (SSSOM)[30] document can define its own prefix map, the SSSOM standard includes a default prefix map (that can, again, be modified or extended) to promote consistency and enable direct downstream data integration. The default prefix map is automatically generated by the Bioregistry with the following settings:

1. The preferred prefix is used instead of the normalized prefix, when available. This is most applicable to OBO Foundry ontologies that typically have a full uppercased preferred prefix (e.g., Chemical Entities of Biomedical Interest (ChEBI)[31]) or a mixed-case preferred prefix (e.g. NCBITaxon).
2. A small set of OBO-influenced remappings of prefixes are applied (`umls` goes to `UMLS`, `snomedct` goes to `SCTID`, and `ensembl` goes to `ENSEMBL`).
3. The URI prefixes used are prioritized to use OBO PURLs, when available.
4. The mapping is restricted to being one-to-one by removing entries with provides and canonical annotations (these fields are described in detail in the data model at https://github.com/biopragmatics/bioregistry/blob/main/docs/datamodel.md), which enables a simple algorithm for URI <-> CURIE conversion based on the prefix map.
5. URI format strings that are not well-behaved (i.e., are not equivalent to a URI prefix) are omitted.

##### 6.2 OBO Foundry

Ontologies can be submitted for consideration for the OBO Foundry through a GitHub issue and are required to contain a certain standard of metadata. However, the standard has changed over time and many entries did not conform. The Bioregistry wrapper

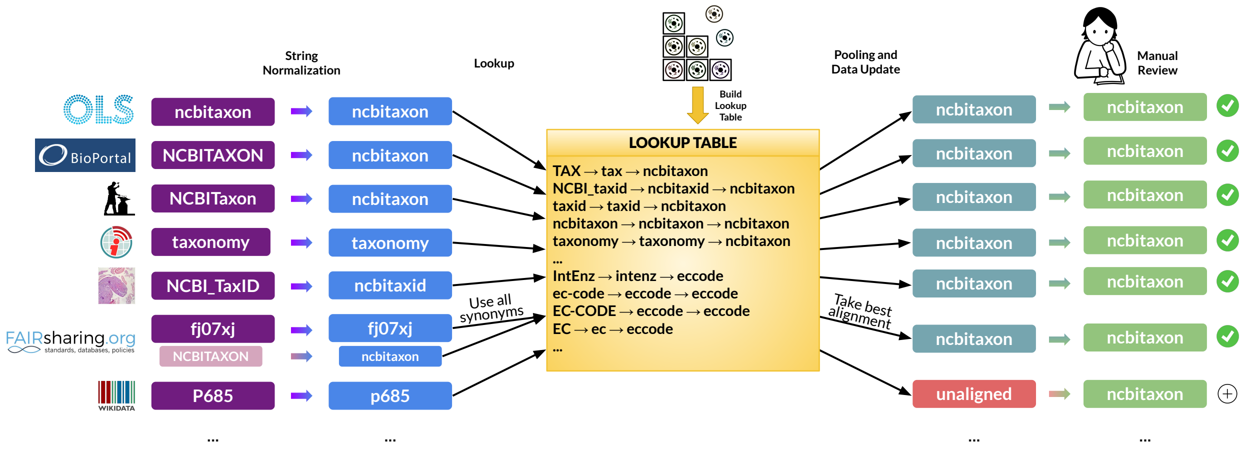


**Figure 2.** A schematic diagram of the automated alignment procedure. First, a lookup table is generated from the prefixes and synonyms curated in the Bioregistry using a standard string normalization workflow (e.g., that maps the synonym `TAX` to `tax` in the context of `ncbitaxon`). Second, external registry records' prefixes (and synonyms, if available) are normalized with the same procedure and checked in the lookup table for a path to a Bioregistry prefix mapping.

of the OBO Foundry metadata made it possible to implement checks and enable directed curation to improve the quality of metadata along several axes (e.g., include trackers in for all ontologies, include repositories for all ontologies, standardize license information for all ontologies, include GitHub contact information for all ontologies) as well as identify some issues that required community involvement (e.g., checking GitHub data for inactive repositories and identify curation errors (e.g., typos) in the Planarian Anatomy Ontology[32] (PLANA).

The Bioregistry's wrapping of the OBO Foundry's metadata has made it accessible and useful for several metadata maintenance scenarios:

1. Implement new guidelines for avoiding prefix conflicts (https://github.com/OBOFoundry/OBOFoundry.github.io/issues/1519)
2. Enable parking prefixes in the Bioregistry so they don't need to be parked in the OBO PURL system (https://github.com/OBOFoundry/OBOFoundry.github.io/issues/1677)
3. Identify and standardize non-standard and undocumented invalid CURIEs throughout OBO Foundry ontologies, in particular database cross-references (https://github.com/obophenotype/planaria-ontology/issues/205)

#### 7 Future Use Cases

This section describes projects that are currently considering or working towards adopting the Bioregistry.

##### 7.1 Biolink Model

Biolink Model[4] is a data schema and metamodel for biomedical data generated with LinkML (https://github.com/linkml). Biolink Model is currently a downstream consumer of Identifiers.org and Prefix Commons identifier. With those services being out of date or slow to respond to new prefix requests (or with the issue above that the desired or historical prefix is a mismatch to the community-established prefix), Biolink Model also encodes its own prefix to URL mapping. A current barrier for this project adopting the Bioregistry is the difficulty of communicating the Bioregistry's features related to prefix capitalization and domain-specific standardization. Several developers and maintainers have expressed interest, but require further education about the differences between the Bioregistry's prefix normalization scheme, its usage of preferred prefixes (implemented in https://github.com/biopragmatics/bioregistry/pull/169 to better support OBO Foundry users), and its ability to generate domain-specific contexts (described in the main text).

##### 7.2 Knowledge Graph Exchange

The Knowledge Graph Exchange (KGX) Python library provides a simplified abstraction for parsing, building, and exchanging knowledge graphs that conform or are aligned to the Biolink Model. The default implementation makes use of networkx.MultiDiGraph for an in-memory representation of a graph but KGX is extendable to other graph libraries. KGX allows users to read from graphs in various formats like RDF serializations, SPARQL endpoints, Neo4j endpoints, CSV/TSV, JSON, OWL, and OBOGraph JSON. KGX also allows users to perform a series of operations on a graph that facilitates an Extract-Transform-Load pipeline. Since KGX is Biolink Model-aware, it can also provide validation on the incoming graphs and write Biolink Model compliant graphs. While KGX (as of v1.5.5) does not use Bioregistry as a dependency, there are a couple of avenues where KGX can benefit from Bioregistry and associated tools. Firstly, KGX has a prefix manager module that can be configured with a JSON-LD context to define prefix to URI mappings. The path of least resistance would be to use the Bioregistry prefix to URI mappings such that incoming URIs are mapped to Bioregistry recommended prefixes and back. Secondly, KGX has methods to manage the transformation of URIs to CURIEs and vice versa, which in turn relies on Prefix Commons. Bioregistry provides several functionalities that KGX can benefit from, for example, the ability to get a normalized prefix for an URI makes it easier to manage the transformation of incoming URIs to standard URIs. Finally, the Bioregistry-normalized prefixes (as provided by bioregistry.context.jsonld) and OBO specific prefixes (as provided by obo.context.jsonld) ensure that frequently encountered prefixes are managed without ambiguity in KGX.

##### 7.3 Alliance Of Genome Resources

Another use case for a single prefix registry can be found in a number of project-specific prefix registries that have been generated independently of a centralized service. For example, the Alliance of Genome Resources[33] uses https://github.com/alliance-genome/agr_schemas/blob/master/resourceDescriptors.yaml to generate links to external resources.

#### 8 Review and Comparision

##### 8.1 Capabilities and Qualities

This section provides a companion to Table 2 of the main text to describe its fields in more detail. A similar explanation is provided at https://bioregistry.io/related#capabilities.

**Structured Data** This field denotes if the registry provides structured access to its data? For example, this can be through an API (e.g., FAIRsharing, OLS) or a bulk download (e.g., OBO Foundry) in a structured file format. A counter-example is a site that must be scraped to acquire its content (e.g, the NCBI GenBank).

**Bulk Data** This field denotes if the registry provides a bulk dump of its data. For example, the OBO Foundry provides its bulk data in a file and Identifiers.org provides its bulk data in an API endpoint. A counterexample is FAIRsharing, which requires slow, expensive pagination through its data. Another counterexample is HL7 which requires manually navigating a form to download its content. While GenBank is not structured, it is still bulk downloadable.

**No Authentication** This field denotes if the registry provides access to its data without an API key. For example, Identifiers.org. As a counter-example, BioPortal requires an API key for access to its structured data.

**Automatable Download** This field denotes if the registry makes its data available downloadable in an automated way. This includes websites that have bulk downloads, paginated API downloads, or even require scraping. A counter example is HL7, whose download can not be automated due to the need to interact with a web form.

**Permissive License** This field denotes if the registry uses a license that permits reuse and or remixing. Based on the OBO Foundry's FP-001 Openness Principle[3], this includes Creative Commons CC BY 3.0, CC BY 4.0, and CC Zero. This explicitly does not include resources licensed with share-alike clauses, no derivatives clauses, or ones that are missing license statements entirely.

**Prefix Search** This field denotes if the registry provides either a dedicated page for searching for prefixes (e.g. AberOWL has a dedicated search page) OR a contextual search (e.g., AgroPortal has a prefix search built in its homepage).

**Prefix Provider** This field denotes if the registry provides information about its own prefixes either in the form of a web page or an API endpoint. These can be accessed through a stable URL into which a prefix from the registry can be formatted.

**CURIE Resolver** This field denotes if the registry can act as a resolver, i.e., it redirects to an external page about a given biomedical concept or entity based on its CURIE and the registry's internal metadata data about the prefix's associated URI format string.

**CURIE Lookup** This field denotes if the registry act as a lookup service, i.e., it gives information about a given biomedical concept or entity based on its CURIE.

[3] https://obofoundry.org/principles/fp-001-open.html
